# Acid exposure impairs mucus secretion and disrupts mucus transport in neonatal piglet airways

**DOI:** 10.1101/669879

**Authors:** Yan Shin J. Liao, Shin Ping Kuan, Maria V. Guevara, Emily N. Collins, Kalina R. Atanasova, Joshua S. Dadural, Kevin Vogt, Veronica Schurmann, Leah R. Reznikov

## Abstract

Tenacious mucus produced by tracheal and bronchial submucosal glands is a defining feature of cystic fibrosis (CF). Although airway acidification occurs early in CF, whether transient acidification is sufficient to initiate mucus abnormalities is unknown. We studied mucus secretion and mucus transport in piglets forty-eight hours following an intra-airway acid challenge. Acid-challenged piglet airways were distinguished by increased mucin 5B (MUC5B) in the submucosal gland but decreased lung lavage fluid MUC5B, following *in vivo* cholinergic stimulation, suggesting a failure in submucosal gland secretion. Concomitantly, intrapulmonary airways were obstructed with glycoprotein rich material under both basal and methacholine-stimulated conditions. To mimic a CF-like environment, we also studied mucus secretion and transport under diminished bicarbonate and chloride transport conditions *ex vivo*. Cholinergic stimulation in acid-challenged piglet airways induced extensive mucus films, greater mucus strand formation, increased dilation of submucosal gland duct openings and decreased mucociliary transport. Finally, to elucidate potential mediators of acid-induced mucus defects, we investigated diminazene aceturate, a small molecule that inhibits the acid-sensing ion channel (ASIC). Diminazene aceturate restored surface MUC5B in acid-challenged piglet airways under basal conditions, mitigated acid-induced airway obstruction, and magnified the number of dilated submucosal gland duct openings. These findings suggest that even transient airway acidification early in life might have profound impacts on mucus secretion and transport properties. Further they highlight diminazene aceturate as an agent that might be beneficial in alleviating certain mucus defects in CF airway disease.

**One sentence summary:** Early life airway acidification has profound impacts on mucus secretion and transport.

## INTRODUCTION

Cystic Fibrosis (CF) is a life-shortening autosomal recessive disorder caused by mutations in the gene encoding the *cystic fibrosis transmembrane conductance regulator (CFTR)*. CF airways are distinguished by frequent infection, inflammation and adherent mucus. Over time, these factors lead to progressive airway destruction and respiratory demise (*1-4*).

Several lines of evidence suggest that airway mucus is abnormal in early CF disease (*5, 6*). For example, pathologic findings in neonates with CF described bronchioles obstruction due to “thick” mucus produced by the tracheal and bronchial submucosal glands (*5*). Similarly, Hoegger and colleagues found a failure of mucus detachment from airway submucosal glands in newborn pigs with CF, resulting in impaired mucociliary transport (*6*). Likewise, rats with CF displayed submucosal gland duct plugging prior to onset of airway infection (*7*), whereas ferrets with CF showed excessive mucus accumulation even in a sterile airway environment (*8*). Combined, these studies highlight the submucosal gland as a key node in CF pathogenesis (*9*), and suggest that abnormal mucus might precede infection and inflammation. Consistent with this, a recent study in preschool children with CF suggested that mucus accumulates in the airways prior to infection and airway remodeling (*10*).

The cause of abnormal mucus in CF has received significant attention (*11*). Numerous mechanisms have been proposed, including dehydration of the airway (*12-15*) and a lack of CFTR-mediated bicarbonate transport (*16*), which acidifies the airway surface (*17, 18*). Both hypotheses are attractive. The lack of bicarbonate transport and an acidic airway pH hypothesis has received scrutiny as studies describing acidification find that it occurs early in disease. For example, decreased airway pH has been found in CF pigs (*18*) and CF rats (*7*) prior to the onset of infection and inflammation, whereas in humans, decreased airway surface pH has been described in neonates with CF (*19*), but not children (*20, 21*).

Our previous data suggested that neonatal piglets challenged with intra-airway acid exhibit airway obstruction (*22*), suggesting defects in mucus secretion, production and/or transport. Here we tested the hypothesis that transient airway acidification is sufficient to induce defects in mucus secretion and mucociliary transport in neonatal piglets. Secondarily, we tested the hypothesis that pharmacological inhibition of acid detection would mitigate acid-induced secretion defects.

## RESULTS

### Airway surface MUC5B is decreased in acid-challenged piglet airways under basal conditions

We performed multi-level analyses of the secretory gel-forming mucins, MUC5AC and MUC5B (*23, 24*). We first examined basal secretion by measuring the amount of MUC5AC and MUC5B in the bronchoalveolar lavage fluid. Detectable levels of proteins were found in all treatment groups. However, no statistically significant differences were noted under basal conditions (Figure 1A, 1B).

**Fig. 1.**
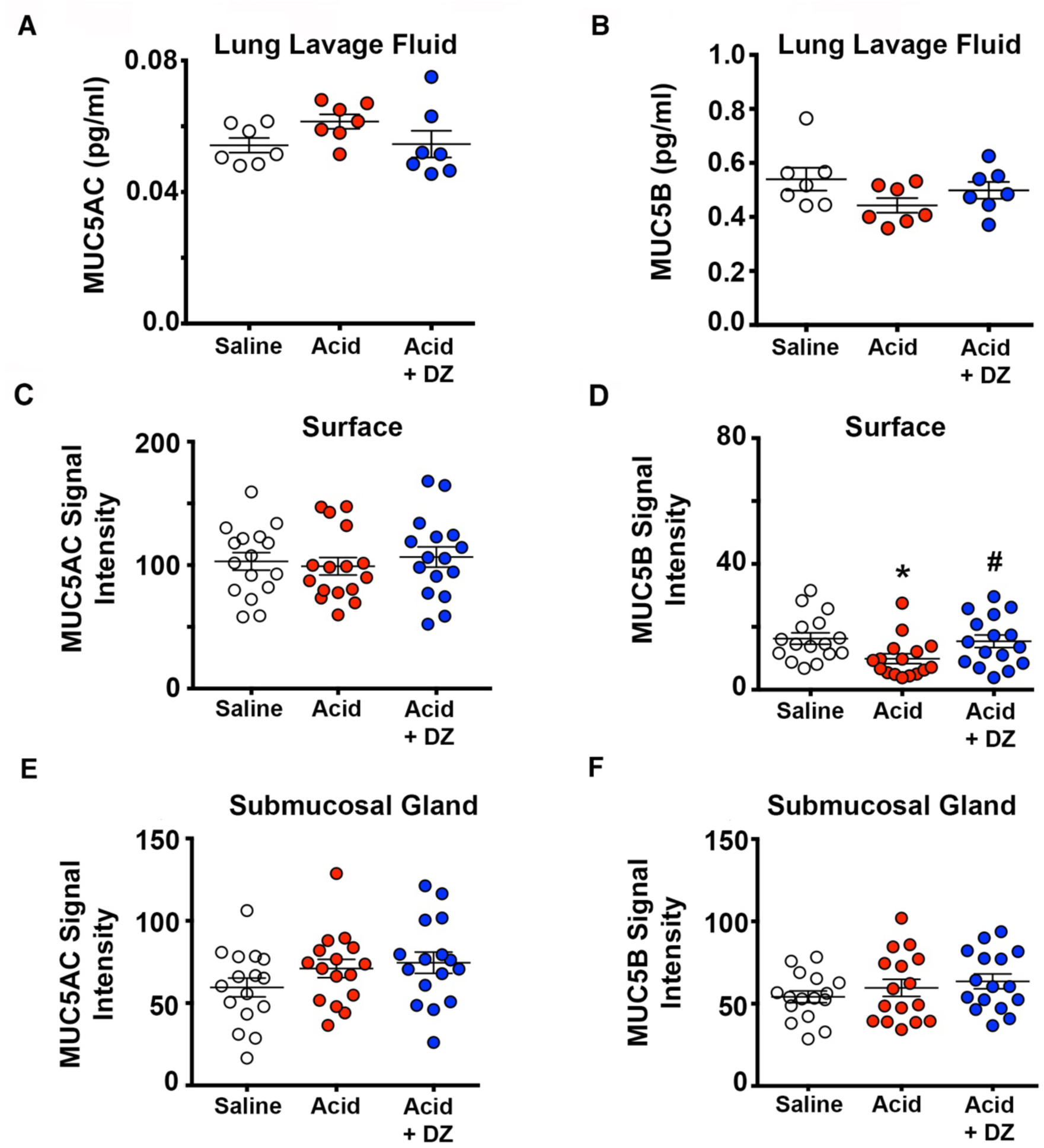
Mucin in piglet airways under basal conditions. Bronchoalveolar lavage fluid concentrations of MU5AC (A) and MUC5B (B). MUC5AC and MUC5B staining signal intensity of the surface epithelia (**C, D**) and submucosal gland (**E, F**). For panels A and B, n = 7 saline-challenged piglets (4 females, 3 males), n = 7 acid-challenged pigs (4 females, 3 males), n = 7 acid-challenged + diminazene aceturate pigs (4 females, 3 males). For panels C-F, n =16 saline-challenged piglets (8 females, 8 males), n = 16 acid-challenged pigs (8 females, 8 males), n = 16 acid-challenged + diminazene aceturate pigs (8 females, 8 males). Data points represent the mean fluorescent intensity for each piglet calculated from 3-5 images analyzed (encompassing the anterior, middle and posterior regions of the trachea). Abbreviations: DZ, diminazene aceturate; MUC5B, mucin 5B; MUC5AC, mucin 5AC. For panel D, * P = 0.031 compared to saline-challenged pigs, # P = 0.036 compared to acid-challenged pigs. Data were assessed with a parametric one-way ANOVA followed by Sidak-Holmes post hoc test. Mean ± S.E.M shown.

Bronchoalveolar lavage fluid is limited in that it primarily captures non-adherent proteins and is a mixed fluid retrieved from alveolar and bronchial spaces. Therefore, we also measured MUC5AC and MUC5B protein expression in tracheal cross sections using antibody-specific labeling and signal intensity analyses (*10*) (Supplemental Figure 1A-C). Analysis of the airway surface revealed no significant differences in MUC5AC signal intensity under basal conditions (Figure 1C). In contrast, acid-challenged piglets had a significant decrease in the signal intensity for airway surface MUC5B compared to saline-treated piglets (Figure 1D). Diminazene aceturate restored MUC5B surface expression in acid-challenged piglets (Figure 1D), suggesting a protective effect.

Decreased surface MUC5B could be due to a decreased basal secretion rate or decreased amount of MUC5B in the submucosal gland or in the airway epithelium. Thus, we also assessed signal intensity for MUC5B and MUC5AC in the submucosal glands but observed no significant differences among treatment groups (Figure 1E, 1F). Quantitative RT-PCR on whole trachea also supported no significant differences in basal expression (Supplemental Figure 2). These data suggested that decreased surface MUC5B was not likely due to a decrease in the amount of MUC5B in the submucosal gland or in the surface epithelium, but highlighted secretion as the likely defect. Further, they suggested that the protective effect of diminizene aceturate was not likely through enhancing production of MUC5B.

### MUC5B secretion is impaired in acid-challenged piglet airways in response to methacholine

If a modest defect in MUC5B secretion under basal conditions was observed, then we reasoned that stimulation of submucosal glands should magnify the defect. Thus, we administered the cholinergic agonist methacholine *in vivo* to a smaller cohort of piglets to stimulate robust submucosal gland secretion. We found no significant differences in MUC5AC concentrations in the bronchoalveolar lavage fluid retrieved from methacholine-stimulated piglets (Figure 2A). In contrast, MUC5B concentrations were significantly decreased in acid-challenged piglets compared to saline-treated controls (Figure 2B). Concentrations of MUC5B in acid-challenged animals provided diminazene aceturate were indistinguishable from acid-challenged piglets (Figure 2B).

**Fig. 2.**
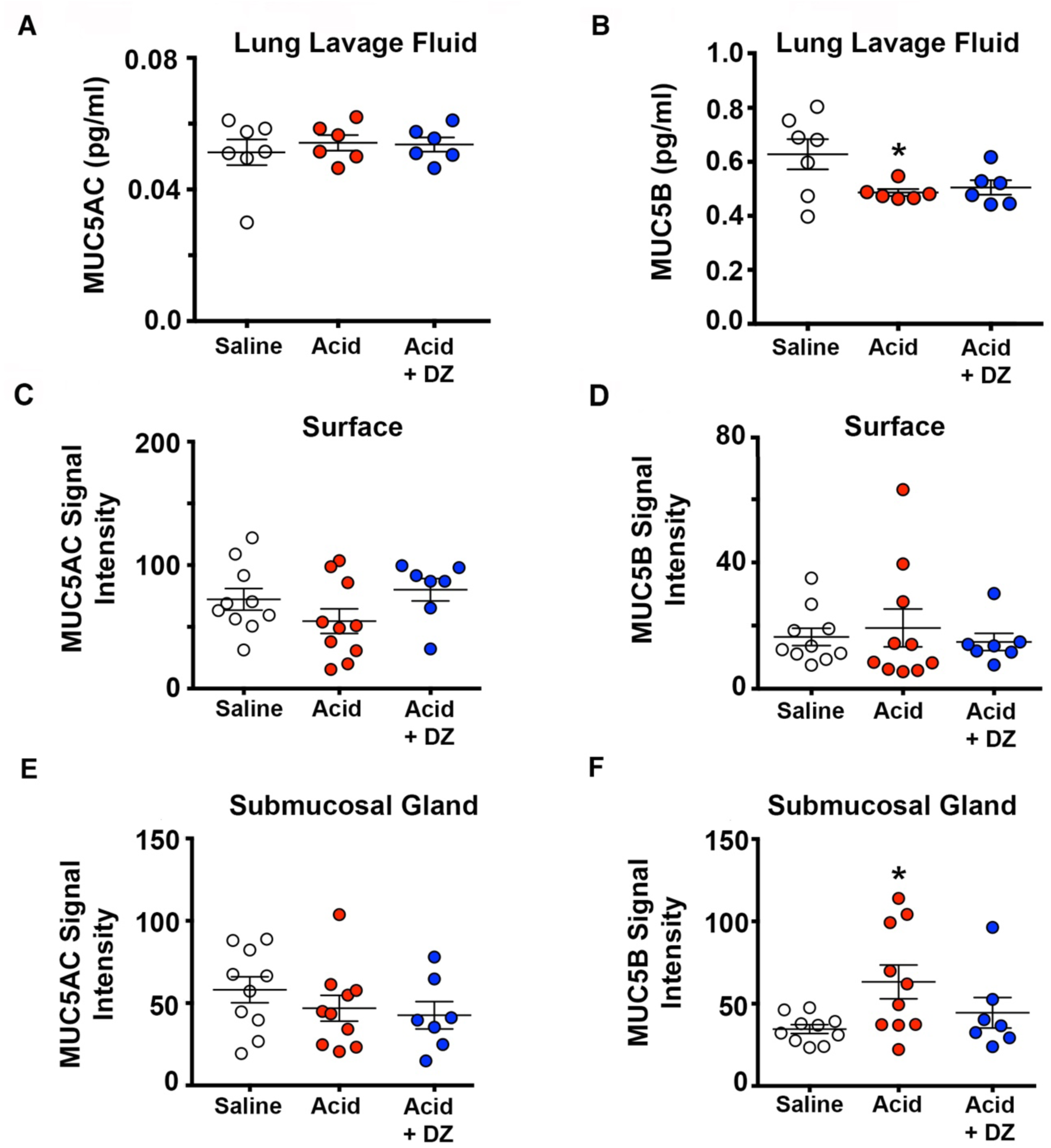
Mucin in piglet airways under methacholine-stimulated conditions. Bronchoalveolar lavage fluid concentrations of MU5AC (**A**) and MUC5B (**B**). MUC5AC and MUC5B staining signal intensity of the surface epithelia (**C, D**) and submucosal gland (**E, F**). For panels A and B, n =10 saline-challenged piglets (5 females, 5 males), n = 10 acid-challenged pigs (4 females, 6 males), n = 7 acid-challenged + diminazene aceturate pigs (4 females, 3 males). For panels C-F, n = 10 saline-challenged piglets (5 females, 5 males), n = 10 acid-challenged pigs (5 females, 5 males), n = 7 acid-challenged + diminazene aceturate pigs (4 females, 3 males). Data points represent the mean fluorescent intensity for each piglet calculated from 3-5 images analyzed (encompassing the anterior, middle and posterior regions of the trachea). Abbreviations: DZ, diminazene aceturate; MUC5B, mucin 5B; MUC5AC, mucin 5AC. For panel B, * P = 0.04 compared to saline-challenged pigs. For panel F, * P = 0.026 compared to saline-challenged pigs. Data were assessed with a parametric one-way ANOVA followed by Sidak-Holmes post hoc test. Mean ± S.E.M shown.

Similar to the basal secretion studies, we also assessed MUC5B and MUC5AC abundance in tracheal cross sections using antibody labeling. No statistically significant differences in the amount of surface MUC5AC (Figure 2C) or MUC5B (Figure 2D) were found. However, when examining the submucosal glands, we found a significant increase in the amount of MUC5B labeling present in acid-challenged piglets compared to saline-treated controls (Figure 2F, Supplemental Figure 3). This finding suggested a retainment of material within the gland, further highlighting a secretion failure. Diminazene aceturate leaned towards significantly mitigating MUC5B secretion defects (Figure 2F, p = 0.12).

### Diminazene aceturate reverts acid-induced mucus obstruction of the small airways

We previously reported that intra-airway acid induces mucus obstruction in the small airways (*22*). Similar to our previous studies, we found that intra-airway acid induced airway obstruction under both basal (Figure 3A-3D) and methacholine-stimulated conditions (Figure 3E-H). Diminazene aceturate significantly attenuated airway obstruction in both conditions (Figure 3C, 3D, 3G, 3H). No differences in the amount of mucin mRNA were noted (Supplemental Figure 4), suggesting that the airway obstruction and alleviation of airway obstruction were not due to differences in mucin expression, but perhaps related to defective secretion. Indeed, mice lacking muc5B show airway obstruction (*25*).

**Fig. 3.**
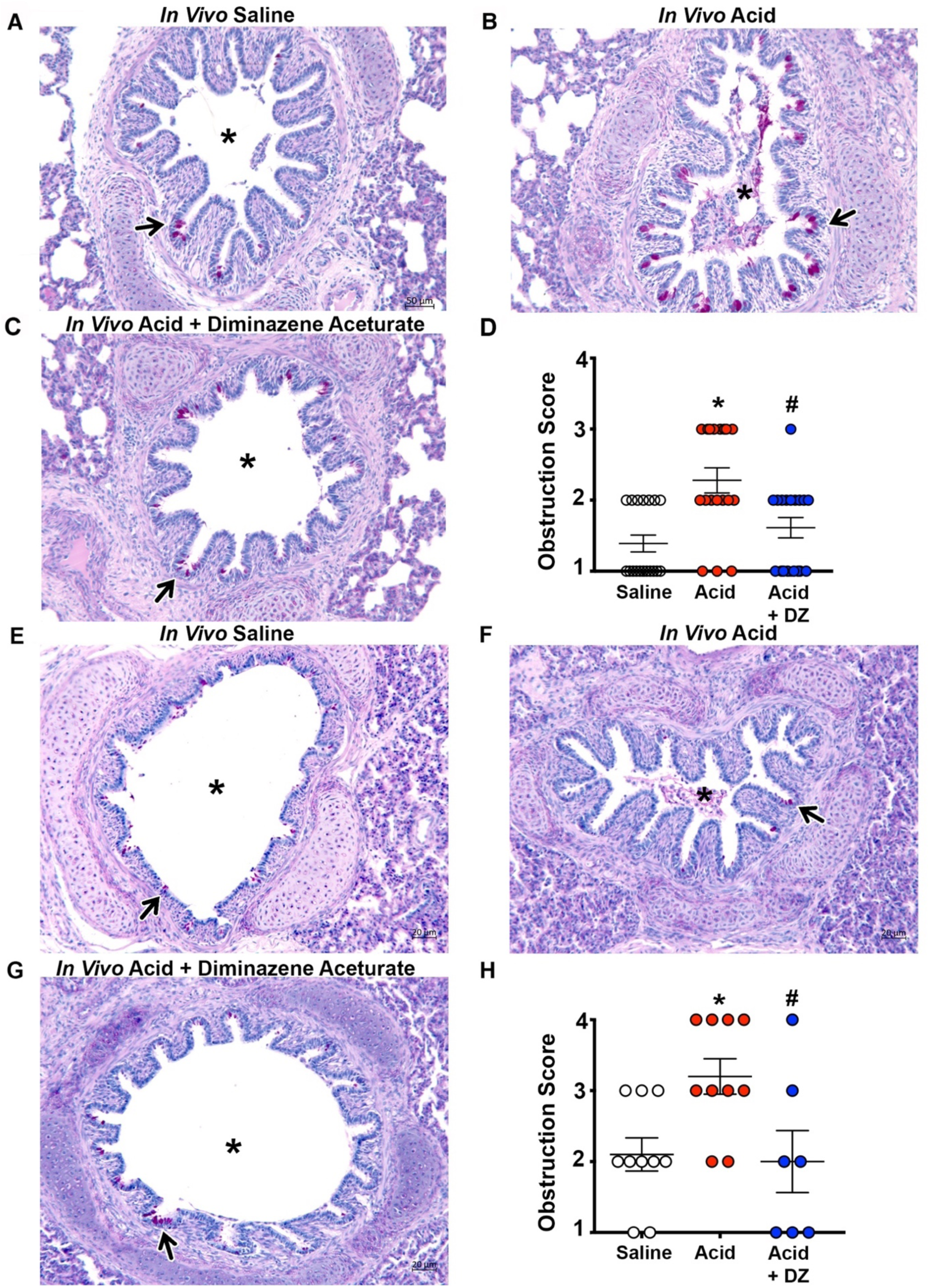
Airway obstruction. Representative histological lung cross sections stained with Periodic acid-Schiff stain (PAS) in piglets challenged with **(A)** saline, **(B)** acid, or **(C)** acid + diminazene aceturate under non-stimulated conditions. Arrows highlight airway and asterisk highlights airway lumen. Scale bar in panel A applies to panels B-C. **(D)** Obstruction score assigned to lung cross sections. n =18 saline-challenged piglets (9 females, 9 males), n = 18 acid-challenged pigs (9 females, 9 males), n = 18 acid-challenged + diminazene aceturate pigs (9 females, 9 males). Representative histological lung cross sections stained with PAS in piglets delivered methacholine in vivo following challenge with **(E)** saline, **(F)** acid, or **(G)** acid + diminazene aceturate. Arrows highlight airway and asterisk highlights airway lumen. Scale bar in panel A applies to panels B-C. **(D)** Obstruction score assigned to lung cross sections. n =10 saline-challenged piglets (5 females, 5 males), n = 10 acid-challenged pigs (4 females, 6 males), n = 7 acid-challenged + diminazene aceturate pigs (4 females, 3 males). Abbreviations: DZ, diminazene aceturate. For panel D, * P = 0.0008 compared to saline-challenged pigs, # P = 0.019 compared to acid-challenged piglets. For panel H, * P = 0.031 compared to saline-challenged pigs, # P = 0.033 compared to acid-challenged piglets. Data were assessed with a non-parametric one-way ANOVA (Kruskal-Wallis) followed by Dunn’s multiple comparison test. Mean ± S.E.M shown.

### Intra-airway acid induces formation of mucus sheets and mucus strands

All of our *in vivo* studies were performed under normal physiological conditions. However, in CF, lack of bicarbonate and chloride transport impacts the morphology and secretion of mucus. Thus, we also investigated mucus morphology and secretion in tracheal segments stimulated with methacholine under diminished bicarbonate and chloride transport conditions. As a surrogate for MUC5AC, we stained mucus with a jacalin lectin (*26*). Wheat germ agglutin (WGA) was used as a surrogate for MUC5B (*26*).

In saline-treated piglet airways, mucus was found in discreet packets that decorated the surface like ornaments (Figure 4A). In contrast, mucus in acid-challenged piglets formed film-like sheets (Figure 4B). Scoring indices indicated greater mucus sheet formation in acid-challenged piglets for jacalin-labeled mucus but not WGA-labeled mucus compared to saline treated controls (Figure 4C, 4D, Supplemental Figure 5). In a smaller cohort of samples, we also assessed sheet formation using IMARIS software. Consistent with our scoring methods, we found a decrease in the number of discreet mucus particles in acid-challenged piglets compared to saline-treated controls (Supplemental Figure 6A). Diminazene aceturate did not significantly alter or prevent the formation of sheet-like structures (Figure 4C, 4D). Finding that acid-challenge augmented the formation of mucus sheets in diminished bicarbonate and chloride transport conditions was consistent with greater accumulation of mucus sheets in CF airways (*26*).

**Fig. 4.**
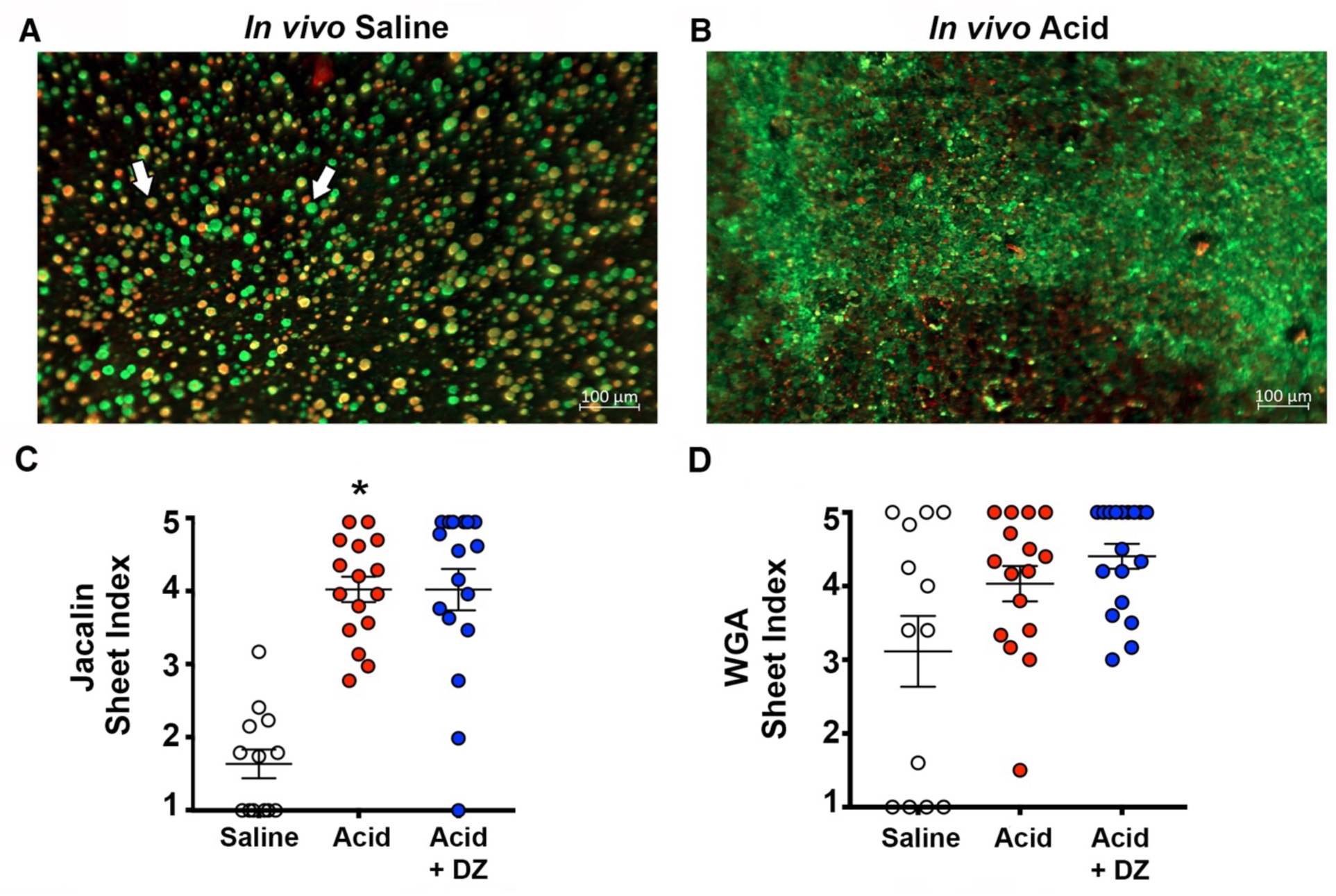
Mucus sheet formation in piglets challenged with intra airway saline, acid, acid + diminazene aceturate under diminished bicarbonate and chloride transport. (**A**) Representative image of an *ex vivo* trachea from a saline-challenged piglet stimulated with methacholine. Discrete entities of mucus were observed and visualized by jacalin lectin (green) and wheat germ agglutinin lectin (red). Arrows highlight examples of lectin labeling. (**B**) Representative image of an *ex vivo* trachea from an acid-challenged piglet stimulated with methacholine. Mucus is visualized by jacalin (green) and wheat germ agglutinin (red) lectins. Sheet index for jacalin-labeled mucus **(C)** and wheat germ agglutinin-labeled mucus **(D)**. n =18 saline-challenged piglets (9 females, 9 males), n = 18 acid-challenged pigs (9 females, 9 males), n = 18 acid-challenged + diminazene aceturate pigs (9 females, 9 males). Data points represent the mean score for each piglet calculated from 5-7 analyzed images (encompassing the anterior, middle and posterior regions of the trachea). Abbreviations: WGA, wheat germ agglutinin; DZ, diminazene aceturate. For panel C, * P = 0.0001 compared to saline-challenged pigs. Data were assessed with a non-parametric one-way ANOVA (Kruskal-Wallis) followed by Dunn’s multiple comparison test. Mean ± S.E.M shown.

Mucus strands (*26*) and bundles (*27*) emanating from submucosal gland ducts have been observed in CF airways. Thus, we examined tracheal segments stimulated *ex vivo* with the methacholine for strand formation (Figure 5, Supplemental Figure 7A-7C). We found no significant differences in the abundance of strands formed by mucus labeled with jacalin (Figure 5C). However, a significant increase in strand formation for WGA-labeled mucus was observed in acid-challenged piglets compared to saline-treated controls (Figure 5D). In a smaller cohort of samples, we performed a secondary analysis that consisted of tracing the strand area manually in image J and expressing it as a percentage of the total image field of view. Consistent with our scoring method, manual tracing also suggested significantly more strands in acid-challenged piglets compared to saline-treated controls (Supplemental Figure 6B). Diminazene aceturate did not significantly alter mucus strand formation (Figure 5C, 5D).

**Fig. 5.**
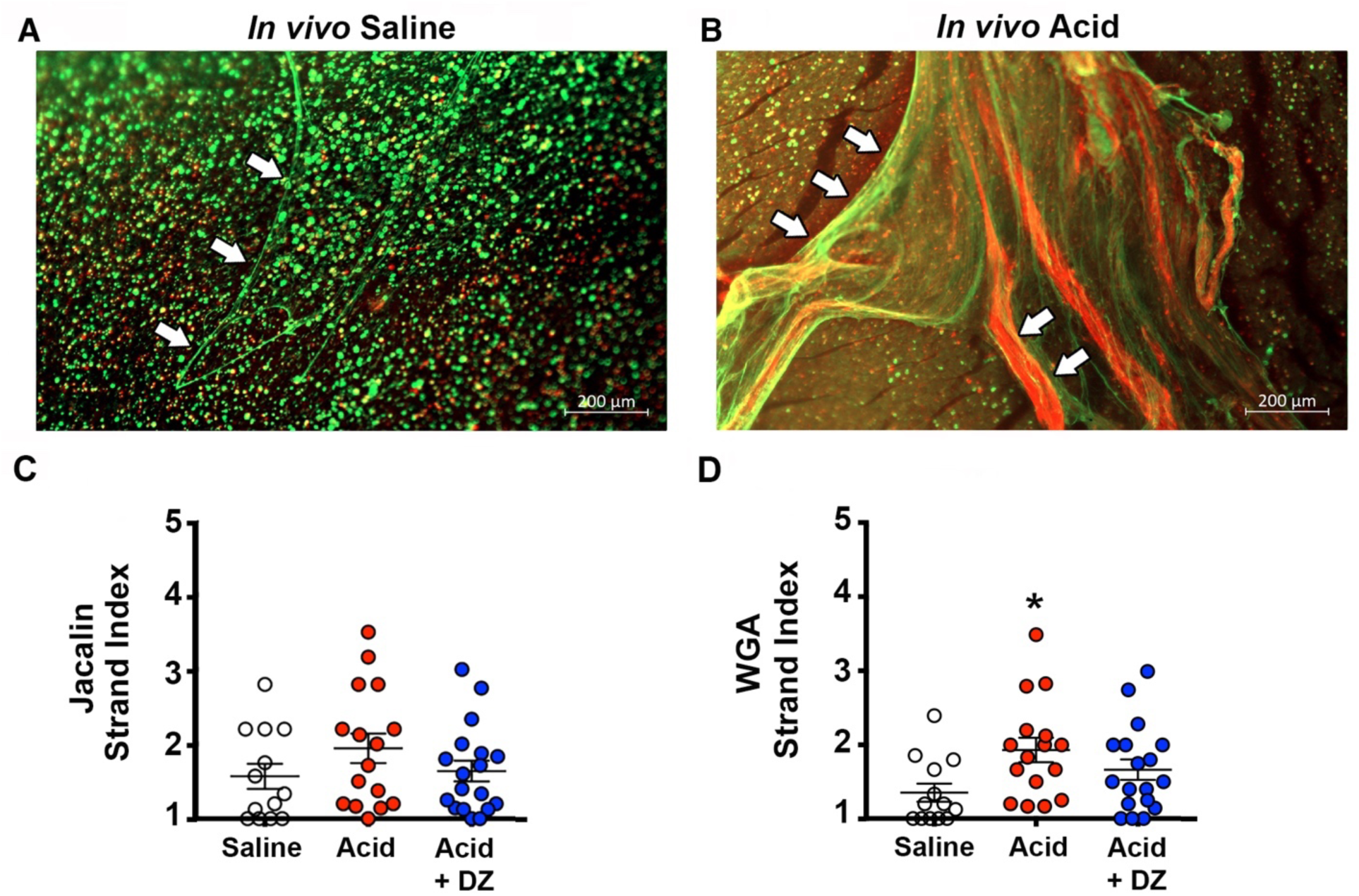
Mucus strand formation in acid-challenged piglets under diminished chloride and bicarbonate transport conditions. (**A**) Representative image of an *ex vivo* trachea from a saline-challenged piglet stimulated with methacholine. Mucus is labeled with jacalin lectin (green) and wheat germ agglutinin lectin (red). Arrows highlight an example of a mucus strand. Representative image of *ex vivo* trachea from a saline-challenged **(A)** and acid-challenged **(B)** piglet airways stimulated with methacholine. Mucus is visualized by jacalin (green) and wheat germ agglutinin (red) lectins. Mucus strands highlighted by white arrows. Strand index for jacalin-labeled mucus **(C)** and wheat germ agglutinin-labeled mucus **(D)**. n = 18 saline-challenged piglets (9 females, 9 males), n = 18 acid-challenged pigs (9 females, 9 males), n = 18 acid-challenged + diminazene aceturate pigs (9 females, 9 males). Data points represent the mean score for each piglet calculated from 5-7 analyzed images (encompassing the anterior, middle and posterior regions of the trachea). Abbreviations: WGA, wheat germ agglutinin; DZ, diminazene aceturate. For panel D, * P = 0.012 compared to saline-challenged pigs. Data were assessed with a non-parametric one-way ANOVA (Kruskal-Wallis) followed by Dunn’s multiple comparison test. Mean ± S.E.M shown.

Dilated submucosal gland ducts openings are commonly observed in CF airways (*5*). Thus, we also examined the tracheal surfaces for the presence of submucosal gland duct openings. Submucosal gland duct openings were only marginally visible in saline-treated piglets (Figure 6A). In stark contrast, acid-challenged piglets displayed a significant elevation in submucosal gland duct openings that appeared dilated (Figure 6B, 6C, Supplemental Figure 7D-7F). Diminazene aceturate magnified the effect of acid and increased the number of dilated submucosal gland openings (Figure 6C).

**Fig.6.**
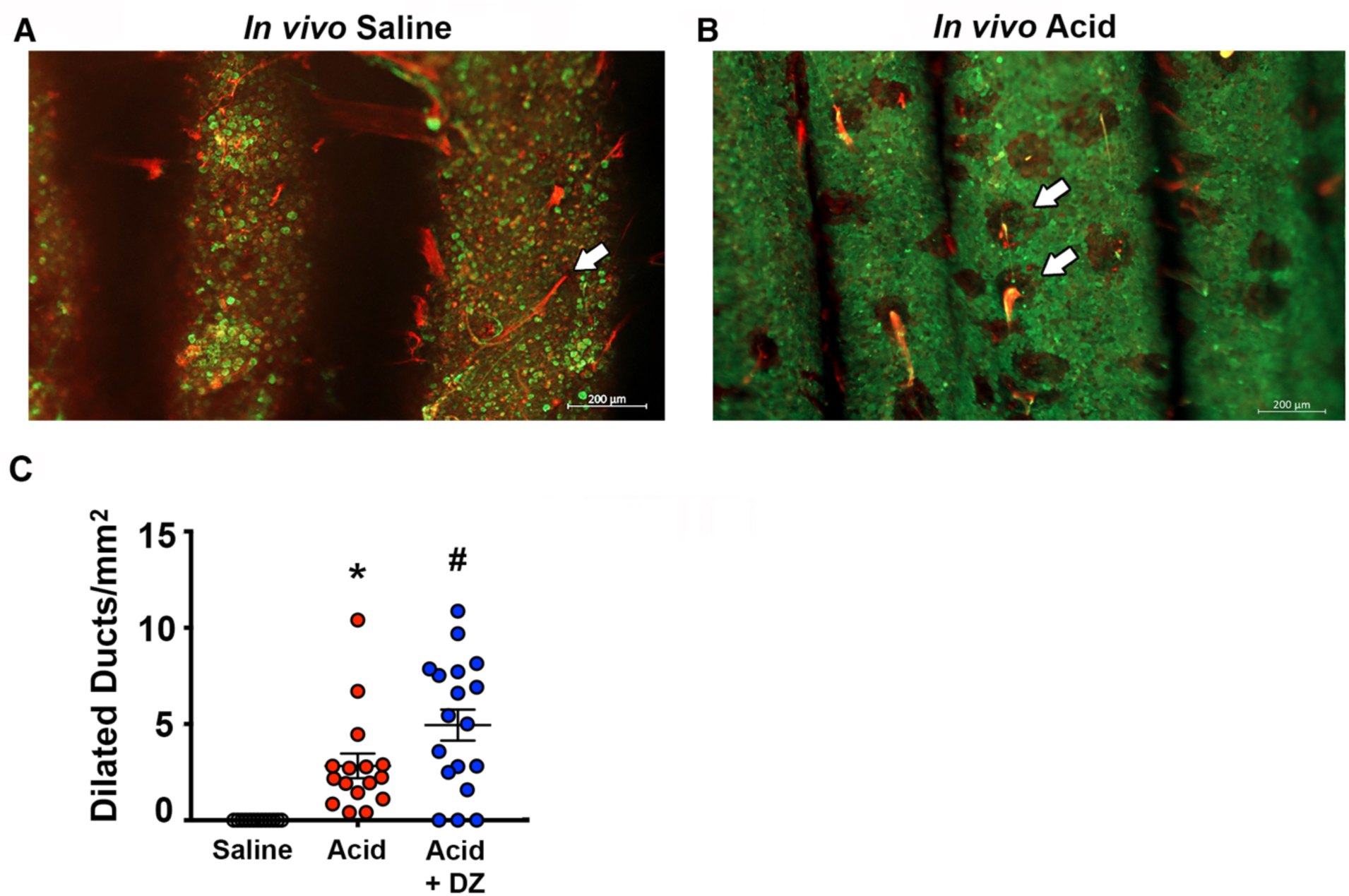
Dilated submucosal gland duct openings on airway surface in acid-challenged piglets. **(A).** Arrows highlight example of a duct opening with mucus emanating in a saline-challenged piglet. **(B)** Arrows highlight examples of a dilated duct openings with mucus emanating in acid-challenged piglets. **(C)** Number of dilated submucosal gland ducts normalized to area. n =18 saline-challenged piglets (9 females, 9 males), n = 18 acid-challenged pigs (9 females, 9 males), n = 18 acid-challenged + diminazene aceturate pigs (9 females, 9 males). Data points represent the mean score for each piglet calculated from 5-7 analyzed images (encompassing the anterior, middle and posterior regions of the trachea). Abbreviations: WGA, wheat germ agglutinin; DZ, diminazene aceturate. For panel C, * P = 0.011 compared to saline-challenged pigs, # P = 0.022 compared to acid-challenged piglets. Data were assessed with a parametric one-way ANOVA followed by Sidak-Holmes post hoc test. Mean ± S.E.M shown.

### Mucociliary transport is impaired in acid-challenged airways

CF airways are distinguished by impaired mucociliary transport (*6, 7, 28, 29*). Thus, to determine whether intra-airway induced mucociliary transport defects, we utilized freshly excised tracheal segments stimulated with methacholine under diminished bicarbonate and chloride transport conditions. For these studies, we utilized methods developed by Hoegger and colleagues (*6*), in which fluorescent nanosphere bind and attach to mucus, allowing for real-time visualization of mucus production and movement. To assess the movement of fluorescently-labeled mucus, we utilized computer assigned particle-tracking (Figure 7A, 7B). We found that both average speed and maximal speed of mucus movement was decreased in acid-challenged piglet airways (Figure 7C, 7D). Because speed is equal to distance over time, we also examined computer assigned particle track length and found it was decreased (Figure 7E). No effect of diminazene aceturate was observed.

**Fig. 7.**
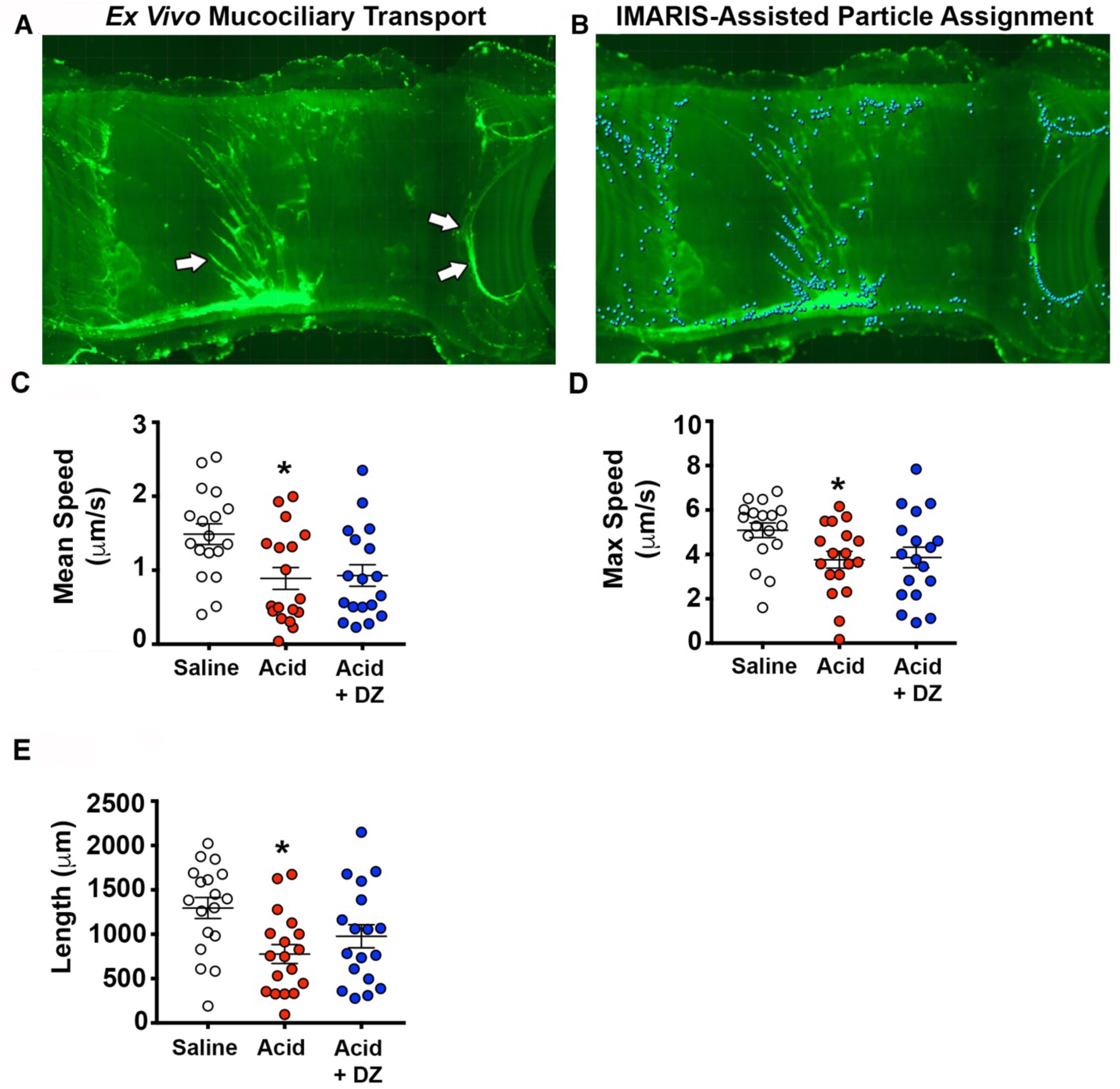
Mucus transport in piglet airways under diminished chloride and bicarbonate transport. **(A)** Representative image of an *ex vivo* piglet trachea stimulated with methacholine. Mucus is visualized in real-time with fluorescent nanospheres (bright green). Mucus often forms strands; examples are highlighted by white arrows. **(B)** Image in panel A that has been processed with Imaris software. Computer particles are assigned based upon fluorescence intensity and appear as blue/aqua in color. Mean mucus transport speed **(C)** and the maximum mucus transport speed **(D). (E)** Computer assigned particle-track length. n =18 saline-challenged piglets (9 females, 9 males), n = 18 acid-challenged pigs (9 females, 9 males), n = 18 acid-challenged + diminazene aceturate pigs (9 females, 9 males). Abbreviations: DZ, diminazene aceturate. For panel C, * P = 0.01 compared to saline-challenged pigs. For panel D, * P = 0.042 compared to saline-challenged pigs. For panel E, * P = 0.007 compared to saline-challenged pigs. Data were assessed with a parametric one-way ANOVA followed by Sidak-Holmes post hoc test. Mean ± S.E.M shown.

## DISCUSSION

In early CF pathogenesis, the airway is acidic (*7, 18, 19*). Although mucus abnormalities precede airway infection and inflammation (*6, 7, 10*), how and whether transient airway acidification impacts the development of mucus abnormalities is unknown. To characterize the significance of acute and transient early life airway acidification on mucus properties, we challenged neonatal piglets with intra-airway acid or saline control. Secondarily, we blocked detection of acid with diminazene aceturate to investigate potential sensors of airway pH (*30*).

Using multi-level approaches, we provided new insight into the consequence of early life airway acidification on mucus secretion and transport properties. Our results showed that under basal conditions, acid-challenged piglet airways showed decreased airway surface MUC5B, suggesting defective submucosal gland secretion. Consistent with that, *in vivo* methacholine stimulation resulted in less MUC5B protein in the bronchoalveolar lavage fluid, and more MUC5B retained in the submucosal gland. Diminazene aceturate restored airway surface MUC5B protein levels under basal conditions but did not prevent the acid-induced defect in methacholine-stimulated secretion.

To mimic a CF-like environment, we investigated mucus secretion and mucus transport under diminished bicarbonate and chloride transport conditions. We observed profound differences in the transport and secretion of mucus. Specifically, greater mucus sheet formation was observed in acid-challenged piglets airways. Additionally, there were a greater number of mucus strands in acid-challenged piglet airways, as well as an elevation the number of submucosal gland duct openings. These features paralleled what has been described in CF pig airways (*26*). Thus, combined, our data suggest that intra-airway acidification greatly impacts mucus secretion and transport properties, inducing many features that mimic CF.

Diminazene aceturate is widely used for the treatment of protozoan diseases and reportedly has no adverse side effects (*31*). Its low cost and availability in numerous regions of the world make it an attractive drug. Diminazene aceturate blocks acid-sensing ion channel 1a (ASIC1a) (*30*). ASIC1a is largely present in nerves innervating the airway (*32, 33*). In our studies, we found a marginal effect of diminazene aceturate in preventing acid-induced mucus defects in the trachea, whereas a strong protective effect of diminazene aceturate was observed in the intrapulmonary airways. It is possible that ASIC1a is more concentrated in the small airways compared to the large airways. It is also possible that ASIC1a is expressed on additional cell types in the small airways compared to the large airways, thus amplifying its protective effect. Further, the degree of acidification might be airway tree-dependent, and thus other sensors in addition to ASIC1a more critical in the larger airways. Surprisingly, we also found that diminazene aceturate magnified the number of submucosal gland duct openings under diminished bicarbonate and chloride transport conditions. However, this was not associated with a greater number of mucus strands, nor a further impairment (or alleviation) of mucus transport defects. Thus, the significance of augmented submucosal gland duct openings induced by diminazene aceturate is unknown and requires additional studies.

Although direct measurements of airway pH in children with CF have shown that pH is not different (*20, 21*), other studies suggest that the airways are acidified in neonates with CF (*19*). These findings have raised debate whether acidification could be an initiating factor in CF pathogenesis. Our studies suggest that even a transient airway acidification early in life is sufficient to induce profound impairments in mucus secretion and transport. These studies therefore highlight that changes in airway pH may be an initiating defect in CF. This is consistent with reports from many CF animal models. For example, in rats with CF, the airways are acidic and submucosal gland duct plugging is observed prior to onset of airway infection (*7*). Hoegger and colleagues found a failure of mucus detachment from airway submucosal glands in newborn pigs with CF, where the airways are also acidic, resulting in impaired mucociliary transport (*6*). Therefore, our studies challenge those that dismiss airway pH as being a critical factor in CF pathogenesis.

We previously examined inflammation extensively in acid-challenged piglets using inflammatory-directed gene arrays and ELISAs, but only found evidence for transcriptional inflammation (*22*). Thus, although it is possible that airway acidification induces mucus abnormalities through local inflammatory-mediated mechanisms, it is unlikely that large-scale and active inflammation is a major driver of acid-induced mucus abnormalities.

Mice lacking MUC5B develop airway obstruction and decreased mucociliary transport (*25*). These features mimic features of CF airway disease. In our studies, acid-challenged piglets showed an impaired MUC5B secretion under basal and methacholine-stimulated conditions. While our findings are consistent with several other studies suggesting defects in submucosal gland secretion (*5-7, 34*), they are slightly at odds with recent data suggesting that CF airways are distinguished by enhanced presence of airway mucins early in life (*10*). However, it is possible that with time, acid-challenged piglets might exhibit increased mucin abundance. It is also possible that inflammation on top of acidification might be required to increase mucin abundance (*10*).

Our study has limitations. We did not assess mucus secretion under basal and stimulated conditions within the same subject. This was largely due to the inability to take airway samples from a subject before and after methacholine stimulation. Further, our study was transient and therefore lacked information regarding long-term consequences of airway acidification. The transient nature of our study, however, might also be an advantage, because it allowed for the effects of acidification to be isolated from potential secondary complications, such as infection and/or prolonged inflammation. Finally, we did not identify a definitive mechanism responsible for acid-mediated defects in mucus secretion and mucus transport, although our data highlight ASIC1a as a contributor. Thus, how acidification induces mucus abnormalities requires additional studies.

In summary, early life airway acidification impaired mucus secretion, evoked mucus obstruction, and decreased mucociliary transport. Diminazene aceturate was partially protective in mitigating acid-induced defects. These findings suggest that even transient airway acidification early in life might have profound impacts on mucus secretion and transport properties. Further, they highlight diminazene aceturate as a potential agent beneficial for alleviating some features of CF airway disease.

## MATERIALS AND METHODS

### Study design

The research objective of this study was to define the impacts of early life airway acidification on mucus secretion and mucus transport. Endpoints measured include mucin protein concentrations in bronchoalveolar lavage fluid, mucin protein expression in tracheal cross-sections, mucin mRNA in whole trachea and lung, airway obstruction in lung samples, mucus secretion and morphology *ex vivo* using lectins, and mucus transport *ex vivo* using live imagine and computer-assisted particle tracking. Subjects were male and female piglets. Data were collected across 15 separate experiments in a controlled laboratory experimental design. Sexes and treatments were balanced within an experiment whenever possible. Male and female piglets were assigned to treatment randomly. Studies were performed by individuals blinded to treatment; only an animal ID was associated with post-mortem samples. Our previous work for histological scoring indicated a “n” of 8 per sex was required to achieve statistical significance (*22, 35*). We observed no sex differences in airway obstruction induced by acid and therefore combined male and female data. Similarly, for qRT-PCR experiments, our previous data indicated that a “n” of 5 was required (*35*), therefore we planned for one additional animal per group per sex. For animals that received methacholine *in vivo*, we performed a smaller study on saline and acid-challenged animals prior to investigating the effects of diminazene aceturate (n = 3 saline-challenged, 3 acid-challenged). For ELISA analysis, we previously reported a n of 6 was required (*22*). Prospective exclusion criteria established included any piglet that required antibiotics or anti-inflammatories due to gastrointestinal illness. Data were not assessed for outliers.

### Animals

A total of 116 piglets (Yorkshire-Landrace, 2-3 days of age) were obtained from a commercial vendor and fed commercial milk replacer (Liqui-Lean) and allowed a 24-hour acclimation period prior to interventions. Data were collected from 15 separate cohorts of piglets across approximately 1-1.5 years. The University of Florida Animal Care and Use Committee approved all procedures. Care was in accordance with federal policies and guidelines.

### Airway Instillation

After acclimation, piglets were anesthetized with 8% sevothesia (Henry Schein). The piglets’ airways were accessed with a laryngoscope; a laryngotracheal atomizer (MADgic) was passed directly beyond the vocal folds as previously described (*22, 35*) to aerosolize either a 500 μl 0.9% saline control or 1% acetic acid in 0.9% saline solution to the airway. This procedure results in widespread distribution of aerosolized solutions throughout the piglet airway, including the lung (*36, 37*).

### Chemicals and Drugs

USP grade acetic acid (Fisher Scientific) was dissolved 0.9% saline to final concentration of 1% and sterilized with a 0.22 µm filter (Millex GP). The pH of the 1% acetic acid solution measured 2.6 using an Accumet AE150 pH probe (Fisher Scientific). The estimated pH once applied to the airway surface is ∼6.6 to 6.8 (*22*). Acetyl-beta-methacholine-chloride (Sigma) was dissolved in 0.9% saline for intravenous delivery and *ex vivo* application. Diminazene aceturate (Selleckchem) was dissolved in 0.9% saline containing 5% DMSO (Fisher Scientific).

### Diminazene aceturate delivery

Approximately 30 minutes prior to airway instillations, piglets were provided an intramuscular injection of either vehicle (5% DMSO + 95% 0.9% saline) or vehicle containing 17.5 mg/ml diminazene aceturate. Vehicle or diminazene aceturate were dosed at 3.5 mg/kg. Dose was chosen based upon previous studies showing efficacy in trypanosome infections in dogs (*38*). Time point was selected based upon previous studies showing peak serum concentrations at 15 minutes and peak interstitial fluid concentrations at 3 hours in adult rabbits (*39*).

### Bronchoalveolar lavage and ELISA

The caudal left lung of each piglet was excised and the main bronchus cannulated; three sequential 5 ml lavages of 0.9% sterile saline were administered as previously described (*22, 35*). The recovered material was pooled, spun at 500 X g, and supernatant removed. A porcine MUC5AC (LSBio, LS-F45847-1) and porcine MUC5B (LSBio, LS-F45852-1) ELISA was performed according to the manufacturer’s instructions. The ELISA was read using a filter-based accuSkan FC micro photometer (Fisher Scientific). The limits of sensitivity were <0.188 ng/ml and < 0.375 ng/ml for MUC5AC and MUC5B, respectively. The intra-assay CV and inter-assay CV were <6.1% and <5.2% for MUC5AC and <6.5% and <5.9% for MUC5B.

### Histology

Lung tissues were fixed in 10% neutral buffered formalin (∼7-10 days), processed, paraffin-embedded, sectioned (∼4 µm) and stained with Periodic-acid-Schiff stain (PAS) to detect glycoproteins as previously described (*22, 35*). Digital images were collected with a Zeiss Axio Zoom V16 microscope. Indices of obstruction were assigned as previously described (*22, 35*).

### RNA isolation and qRT-PCR

RNA from the whole trachea and whole lung were isolated using RNeasy Lipid Tissue kit (Qiagen) with optional DNase digestion (Qiagen). RNA concentrations were assessed using a NanoDrop spectrophotometer (Thermo Fisher Scientific). RNA was reverse transcribed for the whole trachea and whole lung (1000 ng) using Superscript VILO Master Mix (Thermofisher). Briefly, RNA and master mix were incubated for 10 mins at 25°C, followed by 60 mins at 42°C, followed by 5 mins at 85°C. *MUC5AC* and *MUC5B* transcript abundance were measured as previously described (*22, 35*). All qRT-PCR data were acquired using fast SYBR green master mix (Applied Biosystems) and a LightCycler 96 (Roche). Standard ΔΔCT methods were used for analysis (*35*).

### Immunofluorescence in tracheal cross sections

1-2 tracheal rings were removed post-mortem and embedded in Peel-A-Way embedding molds containing Tissue-Tek OCT (Electron Microscopy Sciences). Molds were placed in a container filled with dry ice until frozen and stored at −80 °C long-term. Tissues were sectioned at a thickness of 10μm and mounted onto SuperFrost Plus microscope slides (ThermoFisher Scientific). Storage until immunofluorescence also occurred at −80 °C. We used immunofluorescence procedures similar to those previously described (*22, 40*). Briefly, representative cross-sections from a single cohort of pigs were selected and fixed in 2% paraformaldehyde for 15 minutes. Tissues were then permeabilized in 0.15% Triton X-100, followed by blocking in PBS Superblock (ThermoFisher Scientific) containing 2-4% normal goat serum (Jackson Laboratories). Tissues were incubated with primary antibodies for 2 hours at 37°C. Tissues were washed thoroughly in PBS and incubated in secondary antibodies for 1 hour at room temperature. Tissues were washed and a Hoechst stain performed as previously described (*22*). A 1:10 glycerol/PBS solution was used to cover the sections and cover glass added. Sections were imaged on a Zeiss Axio Zoom V16 microscope. Identical microscope settings within a single cohort of piglet tracheas were used and applied. Three to five images encompassing the posterior, anterior, and lateral surfaces of the trachea were taken. Images were exported and analyzed using ImageJ. The trachea surface epithelia and entire submucosal gland regions were traced and the mean signal intensity MUC5B and MUC5AC recorded. Background signal intensity was measured and subtracted manually. The final signal intensities were averaged to identify a mean signal intensity for MUC5B and MUC5AC per region per each piglet.

### Antibodies and lectins

We used the following anti-mucin antibodies: rabbit anti-MUC5B (1:500; Santa Cruz, Cat.# 20119) and mouse anti-Muc5AC (clone 45M1) (1:1,000, ThermoFisher Scientific, Cat # MA512178), followed by goat anti-rabbit and goat anti-mouse secondary antibodies conjugated to Alexa-Fluor 488 or 568 (ThermoFisher Scientific, 1:1,000 dilution). We used WGA-rhodamine (Vector Laboratories) and Jacalin-FITC (Vector Laboratories) at 1:1,000 dilution. Tracheas were submerged in PBS and visualized using a Zeiss Axio Zoom V16 microscope.

### *In vivo* methacholine and ventilation

We have previously described the ventilation and *in vivo* methacholine procedures in piglets (*22, 35*). Briefly, 48 h post instillation, animals were anesthetized with ketamine (20 mg/kg), and xylazine (2.0 mg/kg), and intravenous propofol (2 mg/kg) (Henry Schein Animal Health). A tracheostomy was performed, and a cuffless endotracheal tube (Coviden, 3.5–4.0 mm OD) was placed. Piglets were connected to a flexiVent system (SCIREQ); paralytic (rocuronium bromide, Novaplus) was administered. Piglets were ventilated at 60 breaths/min at a volume of 10 ml/kg body mass. Increasing doses of methacholine were administered intravenously in approximately ∼3 min intervals in the following doses (in mg/kg): 0.25, 0.5, 1.0, 1.5, 2.0. Piglets were ventilated to ensure animal well-being and patency of airway tissues.

### Mucus secretion assay *ex vivo*

We used methods adopted from Ostedgaard et al (*26*). Briefly, 3-4 rings of trachea were removed post-mortem and the outside of the tracheas were wrapped in gauze soaked with 5 mls of the following: 135 mM NaCl, 2.4 mM K_2_HPO_4_, 0.6 mM KH_2_PO_4_, 1.2 mM CaCl_2_, 1.2 mM MgCl_2_, 10 mM dextrose, 5 mM HEPES, pH 7.4 (NaOH), 1.5 mgs per ml of methacholine, and 100 μM bumetanide. We determined that the amount of methacholine the tissue was exposed to was 0.15 mgs total (e.g., external surface of the trachea is covered by approximately100 μl). Thus, the methacholine dose delivered to the trachea was 0.15mg/4.71cm^2^ (tissue dimensions of trachea: radius = 0.5 cm, height = 1.5 cm). Using a conversion chart for dogs (*41*), we estimated the body surface area of a 2-3 kg piglet to be 0.16-0.21 m^2^. Therefore, the estimated dose of methacholine delivered to the tissue was ∼6-7 fold higher than the cumulative *in vivo* dose of methacholine to account for diffusion of methacholine across multiple tissues layers and in the absence of blood circulation. Tracheas were then placed in a temperature-controlled humidified incubator for 3 hours. Following stimulation, tracheas were fixed overnight in 4% paraformaldehyde and permeabilized for 30 minutes using a triton solution (0.15%), followed by blocking in SuperBlock PBS. Jacalin-FITC and wheat-germ agglutin-rhodamine were used to visual mucus (*26, 42*) and incubated overnight with tracheas at concentration of 1:1,000. Tracheas were then washed, cut upon the posterior surface, and pinned to wax covered petri-dishes followed by submersion in PBS. Tracheas were imaged with Zeiss Axio Zoom V16 posterior to anterior using identical microscope settings. Images were assigned scoring indices for sheet formation and strand formation by two observers blinded to conditions. The number of dilated submucosal gland ducts was also measured by two observers blinded to conditions. Scoring for sheet formation follows: 1 = no sheet formation; 2 = 1-25% of image field shows sheet formation; 3 = 26-50% of image field shows sheet formation; 4 = 51-75% of image field shows sheet formation; 5 = 76-100% of image field shows sheet formation. Sheet formation was defined as the loss of discrete cellular packets of mucus. Secondary analysis and validation of scoring was performed with Imaris software (detailed below). Scoring for strand formation was similar to sheet formation. Scoring for strand formation follows: 1 = no strand formation; 2 = 1-25% of image field shows 3 = 26-50% of image field shows a strand; 4 = 51-75% of image field shows a strand; 5 = 76-100% of image field shows a strand. Secondary analysis consisting of tracing the strand area and expressing it a percentage of the total image field of view was also performed on a subset of images in ImageJ.

### *Ex vivo* mucus transport assays

Mucociliary transport was measured using methods similar to those described by Hoegger et al (*6*). Briefly, 3 rings of tracheas were submerged in 5mls of prewarmed solution containing the following: 138 mM NaCl, 1.5 mM KH_2_PO_4_, 0.9 mM CaCl_2_, 0.5 mM MgCl_2_, 2.67 mM KCl, 8.06 mM Na_2_HPO_4-_7H_2_0, 10 mM HEPES, pH 7.4 (NaOH), and 100 μM bumetanide. Tracheas were placed onto a heated stage and kept at 37°C. Images were acquired every 1 minute for 35 minutes. After 5 minutes of baseline, methacholine was administered directly into the solution covering the basolateral and apical sides of tracheas at a dose of 0.004 mg/ml. This dose matched the estimated accumulative dose of methacholine administered to piglets *in vivo (e.g., 3.75 mg/kg = 3.75mg/L = 0.00375mg/ml).* Mucus transport was assessed for an additional 30 minutes. Imaris software was used to track mucus transport across time. Details about Imaris software and processing are highlighted below.

### Imaris computer-assigned particles for measurement of mucus transport and mucus sheet formation

Computer particles based upon signal intensity above background were automatically generated with Imaris software (Bitplane). The particles were then tracked through time using a custom Imaris algorithm utilized principles of the well-validated algorithms published by Jaqaman and colleagues (*43*). For each trachea, the average mean speeds and max speeds of the computer-assigned particles were reported. The length of the particle track was also computed automatically and the mean track length of all the particles per trachea were calculated and reported. For determination of sheet formation, an image was chosen at random from a subset of tracheas stained with lectins. Particles were assigned to the images automatically as described above. The number of particles (representing the jacalin-labeled mucus with intensity above background) were reported. In tracheas that exhibit robust mucus sheet formation, the number of computer-assigned particles is decreased because of the significant jacalin labeled mucus that covers the field of view (e.g., less signal to noise), as opposed to the tracheas in which there is discreet packets of jacalin-labeled mucus (e.g., greater signal to noise).

### Statistical analysis

For parametric data that compared three groups, we used a one-way ANOVA followed by a Sidak-Holmes multiple comparison test. For non-parametric data that compared three groups, we used a one-way ANOVA (Kruskal-Wallis) test followed by Dunn’s multiple comparison test. For analyses that compared two groups, we used a two-tailed unpaired Students t-test. All tests were carried out using GraphPad Prism 7.0a. Statistical significance was determined as P < 0.05.

## List of Supplementary Materials

### Acknowledgement

We thank Dr. Dave Meyerholz and Dr. Mark Hoegger for helpful comments and suggestions. The authors also thank Dr. Igancio Aguirre for providing technical support and resources.

## Funding

This work was funded by R00HL119560 (PI, LRR) and 10T2TR001983 (Co-I, LRR).

## Author contributions

L.R.R, YS.J.L, SP.K, M.V.G, E.N.C and K.R.A participated in the conception and design of the research. L.R.R, SP.K, YS.J.L, J.S.D, E.N.C, M.V.G, V.S and K.V performed the experiments. L.R.R, SP.K, YS.J.L, E.N.C, M.V.G, and K.R.A analyzed the data. L.R.R, YS.J.L, SP.K, J.S.D, E.N.C, M.V.G, K.R.A interpreted the results of the experiments. L.R.R and K.R.A prepared the figures. L.R.R and M.V.G. drafted the manuscript. All authors edited and reviewed the manuscript.

## Competing interests

The authors declare no competing interests.

## Data and materials availability

All data is available in the manuscript or the supplementary materials.

---

**Fig. S1.**
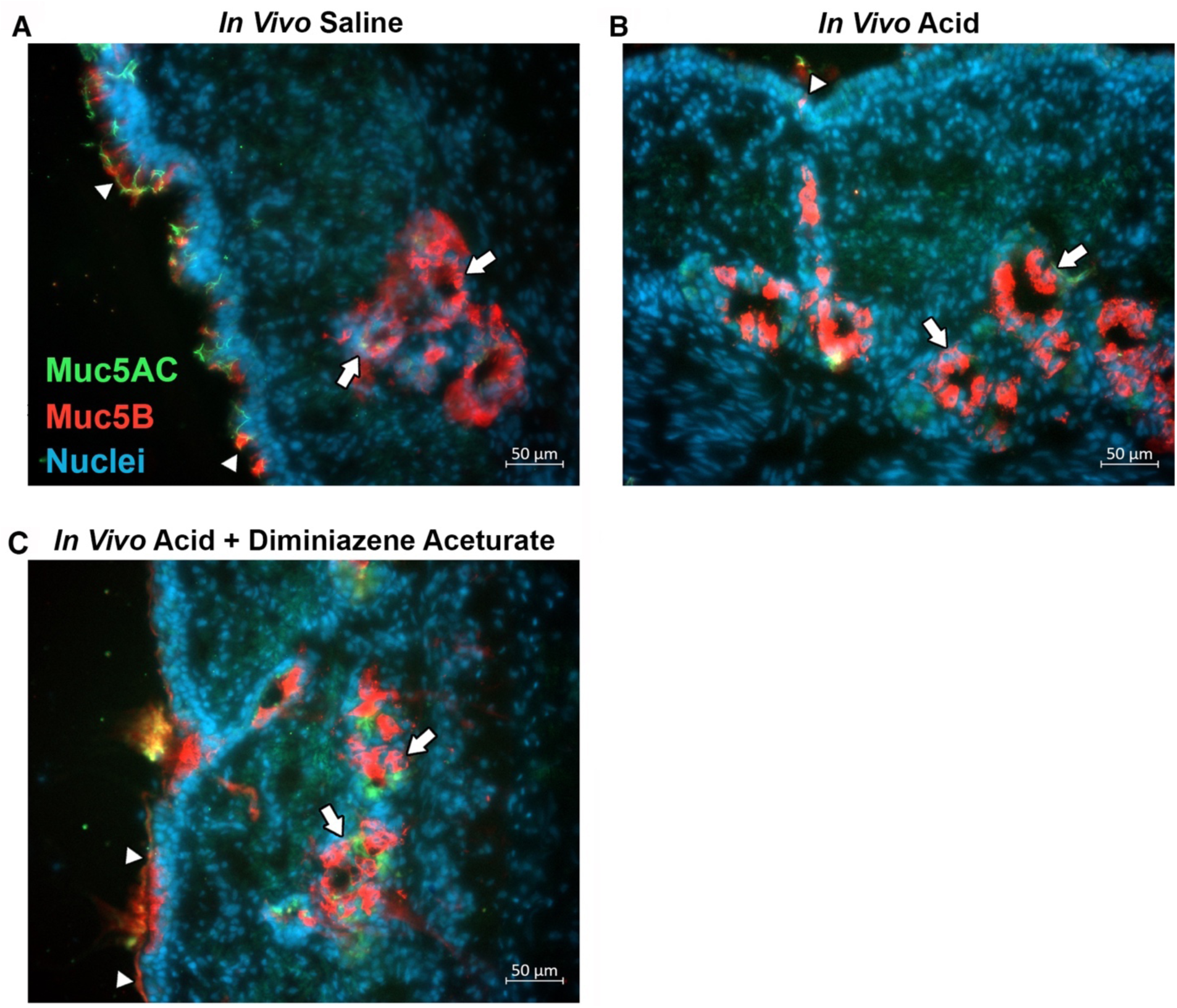
Mucin staining in piglet tracheal cross-sections under non-stimulated conditions. Representative antibody-labeling of mucin 5AC (green) and mucin 5B (red) in tracheal cross sections from piglets challenged with intra-airway saline **(A)**, intra-airway acid **(B)**, intra-airway acid + diminazene aceturate **(C).** Arrow heads represent labeling of mucus on the surface; arrows represent labeling within submucosal glands. Nuclei are shown in blue. Abbreviations: muc5B, mucin 5B; muc5AC, mucin 5AC.

**Fig. S2.**
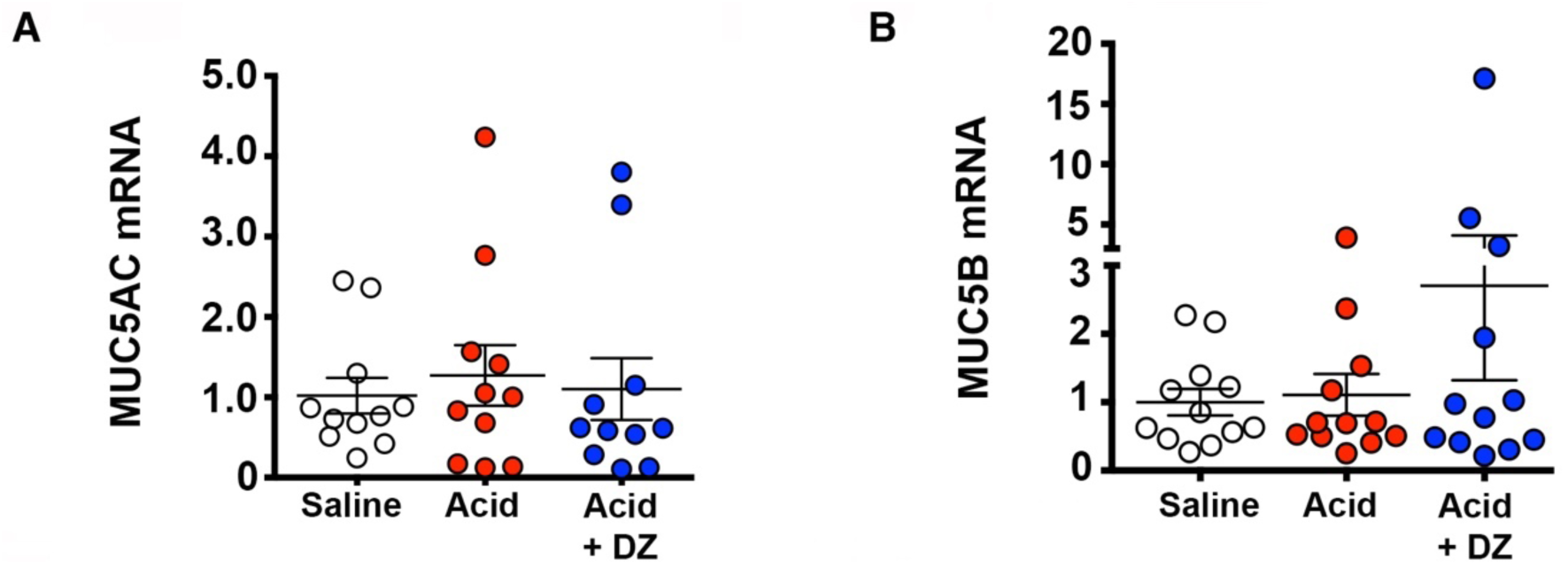
MUC5AC and MUC5B mRNA in whole trachea. mRNA expression for MUC5AC **(A)** and MUC5B (B) in whole trachea. Data are relative to saline-treated controls. n =11 saline-challenged piglets (5 females, 6 males), n = 11 acid-challenged pigs (5 females, 6 males), n = 11 acid-challenged + diminazene aceturate pigs (5 females, 6 males). Abbreviations: DZ, diminazene aceturate; muc5B, mucin 5B; muc5AC, mucin 5AC. Mean ± S.E.M shown.

**Fig. S3.**
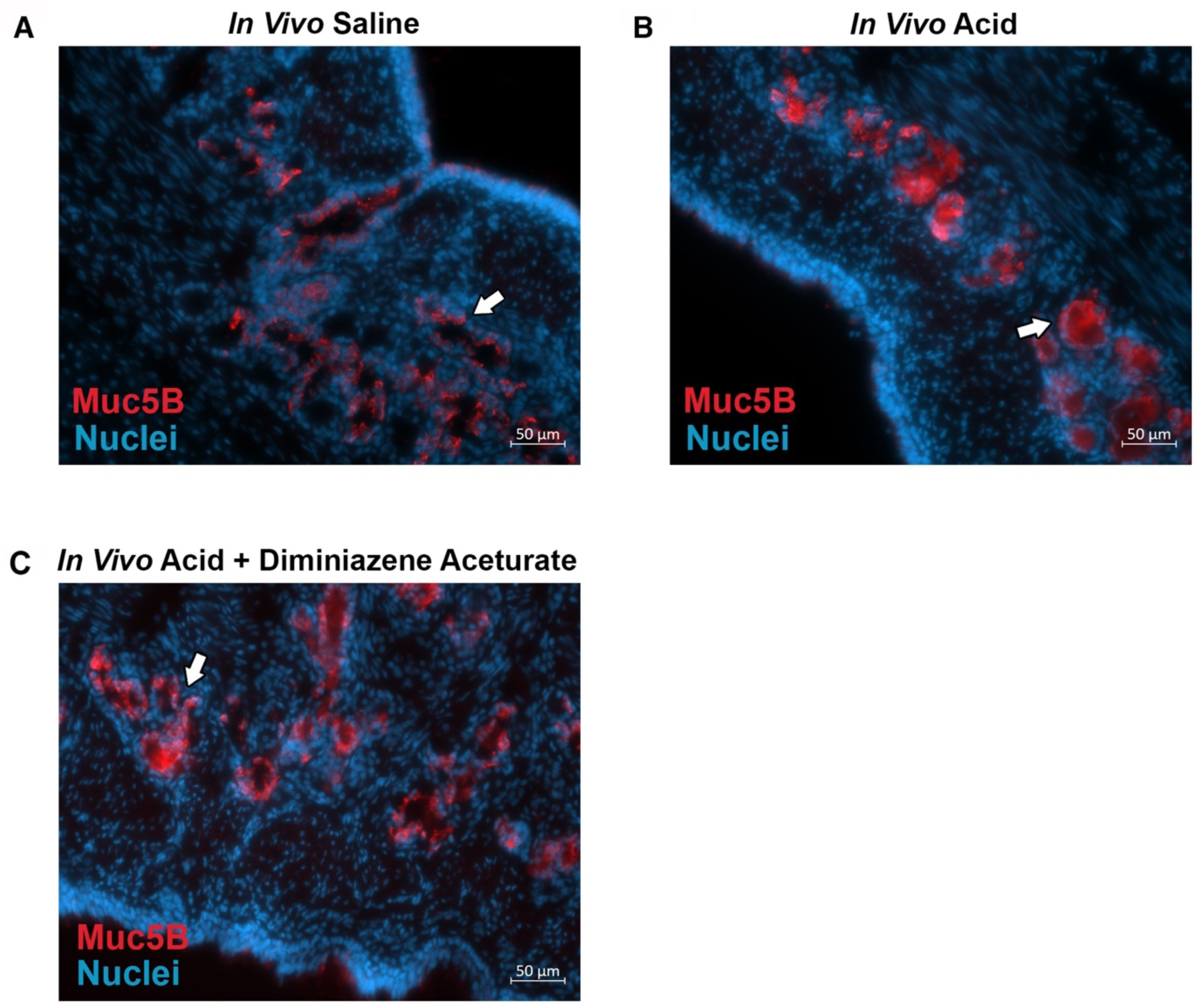
MUC5B staining in piglet tracheal cross-sections following methacholine-stimulation in vivo. Representative antibody-labeling of MUC5B (red) in tracheal cross sections from piglets challenged with intra-airway saline **(A)**, intra-airway acid **(B)**, intra-airway acid + diminazene aceturate **(C).** Arrows represent labeling within submucosal glands. Nuclei are shown in blue. Abbreviations: muc5B, mucin 5B.

**Fig. S4.**
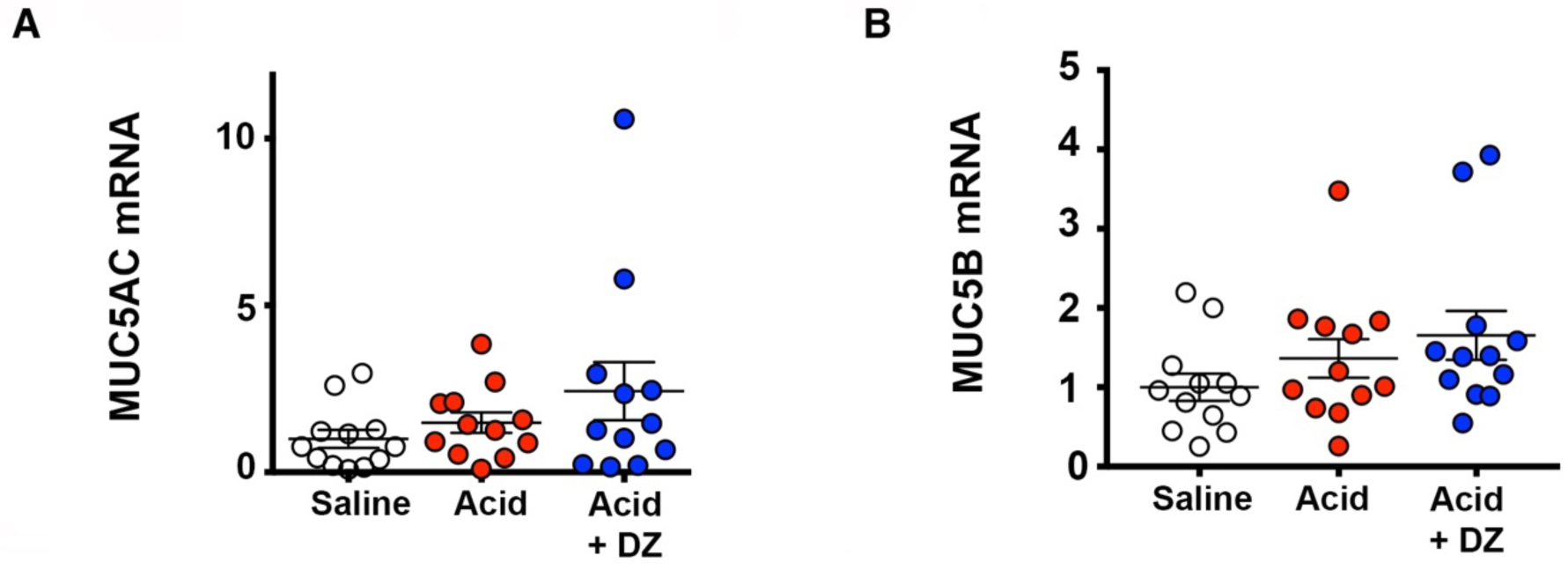
MUC5AC and MUC5B mRNA in whole lung. mRNA expression for MUC5AC **(A)** and MUC5B (B) in whole lung. Data are relative to saline-treated controls. n =12 saline-challenged piglets (6 females, 6 males), n = 12 acid-challenged pigs (6 females, 6 males), n = 12 acid-challenged + diminazene aceturate pigs (6 females, 6 males). Abbreviations: DZ, diminazene aceturate; muc5B, mucin 5B; muc5AC, mucin 5AC. Mean ± S.E.M shown.

**Fig. S5.**
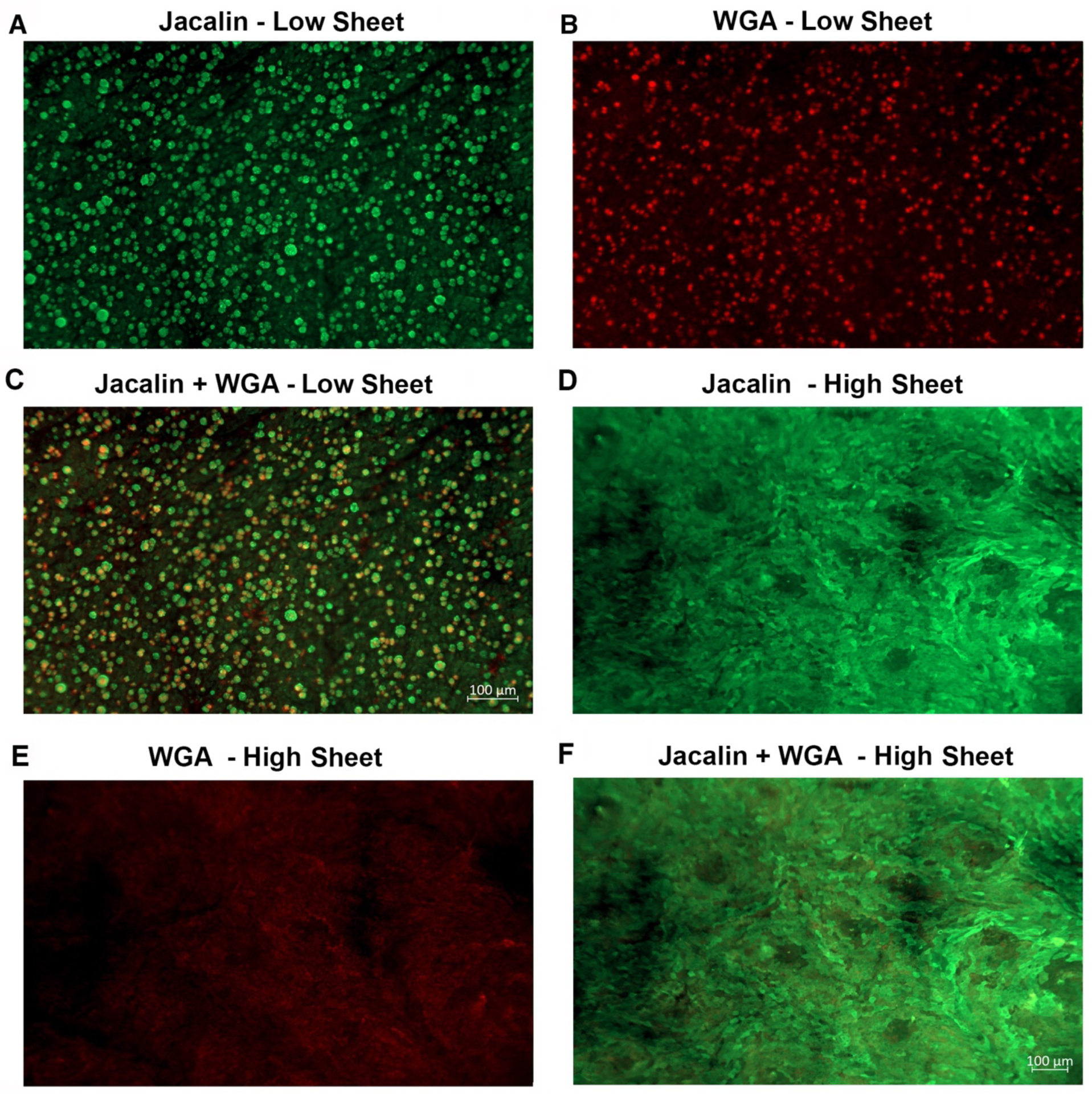
Low and high mucus sheet formation in piglet airways. Representative image of a low sheet formation trachea stained with jacalin and wheat germ agglutinin lectins **(A-C)**. Discrete cellular entities of mucus are observed and illustrated by arrows. Scale bar in panel C applies to panels A and B. Representative image of a high sheet formation in trachea stained with jacalin and wheat germ agglutinin lectins **(D-F)**. Note loss of discreet cellular packets of mucus. Scale bar in panel F applies to panels D and E. Abbreviations: WGA, wheat germ agglutinin.

**Fig. S6.**
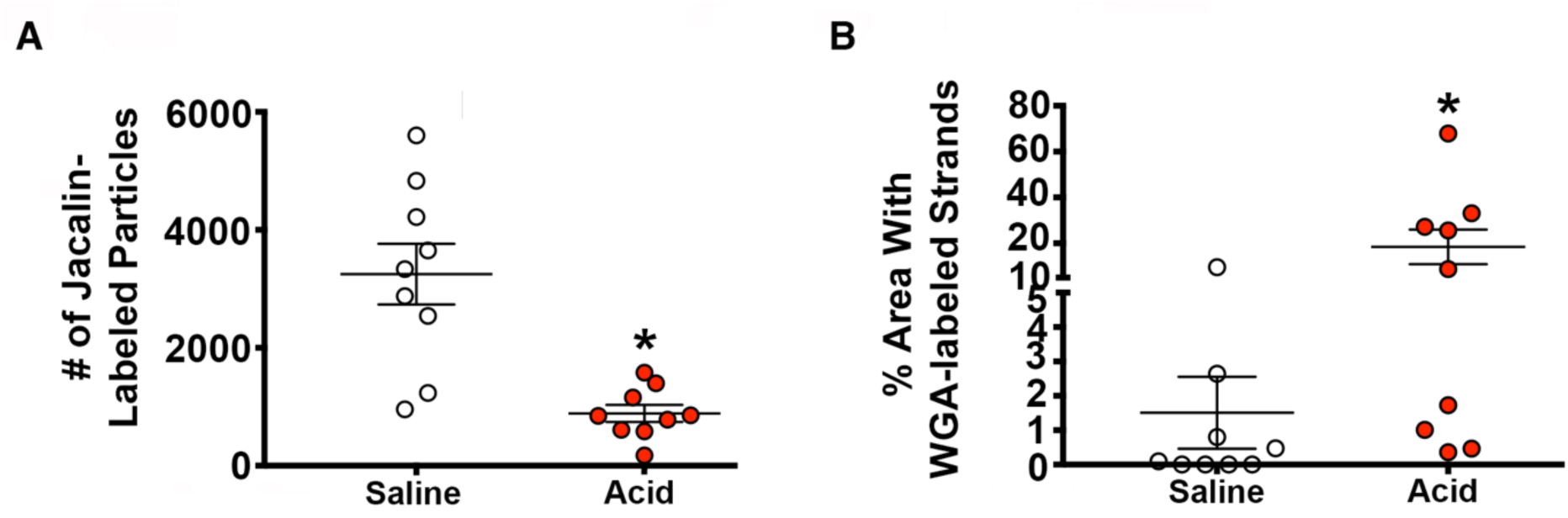
Jacalin-labeled mucus particles on trachea surface and WGA strand formation. **(A)** The numbers of jacalin-labeled mucus particles were analyzed using IMARIS software as an additional measurement of mucus sheet formation in acid-challenged piglets. n = 9 saline-challenged piglets (4 females, 5 males), n = 9 acid-challenged pigs (4 females, 5 males). **(B)** The percent (%) area of field of view occupied by WGA-labeled strands). n = 9 saline-challenged piglets (4 females, 5 males), n = 9 acid-challenged pigs (4 females, 5 males). Abbreviations: WGA, wheat germ agglutinin. For panel A, * P = 0.0004 compared to saline-challenged pigs. For panel B, * P = 0.041 compared to saline-challenged pigs. Data were assessed with a two-tailed unpaired Students t-test. Mean ± S.E.M shown.

**Fig. S7.**
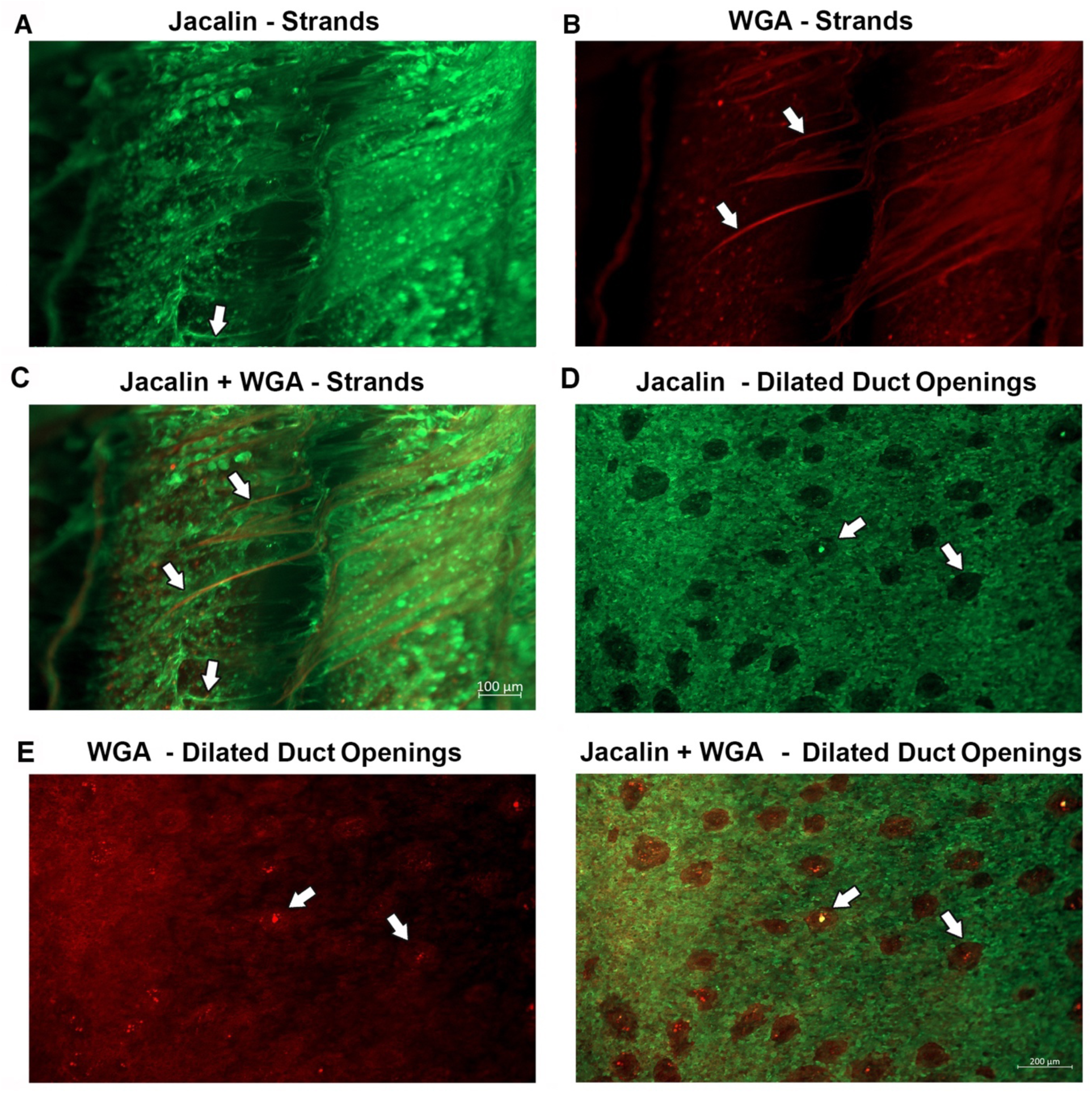
Mucus strand formation and dilated submucosal gland ducts in piglets. Representative image mucus strand formation as detected by jacalin (green) and wheat germ agglutinin (red) lectins **(A-C)**. Arrows highlight an example of a mucus strand for each lectin. Scale bar in panel C applies to panels A and B. Representative image of dilated submucosal gland ducts in a trachea stimulated with methacholine *ex vivo* **(D-F).** Jacalin (green) and wheat germ agglutinin (red) lectins are used to visualize mucus. Arrows highlight examples of dilated submucosal gland duct openings for each lectin. Scale bar in panel F applies to panels D and E. Abbreviations: WGA, wheat germ agglutinin.

